# Lettuce seedlings rapidly assemble their microbiome from the environment through deterministic processes

**DOI:** 10.1101/2024.01.05.574372

**Authors:** Nesma Zakaria Mohamed, Leonardo Schena, Antonino Malacrinò

**Author notes:** corresponding authors: (AM); (LS).

## Abstract

Plant-associated microorganisms have significant impacts on plant biology, ecology, and evolution. Although several studies have examined the factors driving variations in plant microbiomes, the mechanisms underlying the assembly of the plant microbiome are still poorly understood. In this study, we used gnotobiotic plants to test (i) whether seedlings create a selective environment and drive the assembly of root and leaf microbiomes through deterministic or stochastic processes, and (ii) whether seedlings structure the microbiome that is transferred through seeds using deterministic processes and whether this pattern changes when seedlings are exposed to the environmental microbiome. Our results show that the microbiome of gnotobiotic plants (i.e., inherited through seeds) is not under the selective influence of the host plant but changes quickly when plants are exposed to soil microbiomes. Within one week, plants were able to select microorganisms from the inocula, assemble the root microbiome, and assemble the shoot microbiome. This study supports the hypothesis that plants at early developmental stages might exert strong selective activity on their microbiomes and contribute to clarifying the mechanisms of plant microbiome assembly.

## Introduction

Plants grow in close association with a large and diverse community of microorganisms (e.g., bacteria, fungi, nematodes, and viruses) that have profound effects on plant biology, ecology, and evolution [1]. Indeed, the plant microbiome can influence a multitude of host traits, including fitness, nutrient/water uptake, and resistance to biotic and abiotic stressors [1–3]. The structure of the plant microbiome is highly variable throughout developmental stages, across and within plant organs, and between species and genotypes, and is influenced by the physiological status of the plant [1]. While several studies have focused on describing the variation in microbiomes between different plants (e.g., between genotypes), within the same plant (e.g., between organs), or on inferring the effects of different factors (e.g., water, herbivory, pathogens, and agricultural practices) on the structure of the plant microbiome, little is known about the processes that drive the assembly of the plant microbiome. Understanding the mechanisms behind plant microbiome assembly is crucial for leveraging the power of plant-microbe interactions for sustainable agriculture [4].

Plants acquire their microbiome either horizontally from the environment (e.g., soil and air) or vertically from seeds [5–7]. Previous research has shown that soil is a major source of the plant microbiome, whereas the inherited microbiome has a smaller influence [1, 5]. Deterministic and stochastic processes are the major forces driving the assembly of plant microbiomes [8]. Deterministic processes (i.e., selection) influence the presence/absence and abundance of microbial taxa, and are driven by selective forces generated by the host plant or abiotic environment. Deterministic processes can generate dissimilar (variable selection) or similar (homogeneous selection) microbial communities. On the other hand, stochasticity (e.g., dispersal and drift) dominates when selection is weak, and non-selective processes are mainly responsible for driving the assembly of the plant microbiome, such as movements between communities (dispersal) and changes in population size due to random events (drift). Previous studies have focused on testing whether deterministic or stochastic processes dominate the assembly of the plant microbiome; however, results from previous studies have produced contrasting outcomes. There is evidence of deterministic processes driving the assembly of leaf and root microbiomes [9–15], but also evidence suggesting that stochastic processes are dominant [16–21]. Some studies suggest that the assembly of bacterial and fungal communities is often driven by contrasting processes, which may vary according to the plant organ [9, 14, 22, 23]. Other studies have suggested that the dominance of either deterministic or stochastic processes varies over time [14, 24] or as an effect of stress [19, 25]. Thus, current evidence does not show clear patterns in the processes driving the plant microbiome across different plant species.

Previous research has mainly focused on plants grown under field conditions that are already in an advanced stage of growth. This might not provide a complete picture of the dynamics behind the processes that drive plant microbiome assembly. For example, the contribution of deterministic and stochastic factors might change with plant development [14, 24, 26] or plants at a later growth stage might exert a lower level of selection on their microbiome and direct resources to other tasks. Little is known about the processes driving microbiome assembly in plants during the early growth stages. Previously, it was suggested that the assembly of seedling-associated microbial communities might be highly subjected to priority effects and thus are mainly shaped by stochastic processes [26]. However, a few studies have shown that the plant microbiome at the early stages is assembled in a selective environment [14, 22], which might be driven by plants [27]. Indeed, creating a selective environment and directing microbiome assembly processes might be more important for plants at early stages of growth, as this might help them gather beneficial microorganisms to aid plant nutrition and protection against pathogens.

In this study, we used gnotobiotic plants to gain further understanding of the processes driving the assembly of plant microbiomes, particularly immediately after germination, as this is a crucial step when plant-microbiome interactions are established. Lettuce plants (*Lactuca sativa* L.) were grown under gnotobiotic conditions and exposed to 21 different soil microbial communities. After one week, we collected samples (roots and shoots) for amplicon metagenomic (16S and ITS) analyses. First, we investigated whether seedlings create a selective environment belowground and, through selection, drive the assembly of root and leaf microbiomes through deterministic processes. We hypothesized that seedlings create a selective environment belowground, and that through deterministic processes, they assemble both root and shoot microbiomes. Second, we tested whether seedlings exert the same selective forces on the microbiome that is transferred through seeds, or whether selective forces come into play when seedlings are exposed to the environmental microbiome. We hypothesized that the inherited seed microbiome is not subjected to further selective forces [5], but selection occurs quickly once plants are exposed to complex soil microbial communities.

## Results

Amplicon metagenomic sequencing yielded 9,291,816 raw reads for 16S rRNA and 6,904,536 raw reads for ITS. After cleanup and removal of plastidial reads, the 16S dataset included 964,167 reads (average 14,608 reads/sample; min 1,009; max 41,018; Fig. S1A, Fig. S2), whereas the ITS dataset included 1,570,286 reads (average 26,171 reads/sample; min 1,244; max 76,034; Fig. S1B, Fig. S3).

First, we tested whether the plant microbiome (roots and shoots) differed from the composition of the inoculated microbial communities. We found that the multivariate structure of microbial communities differed among inocula, shoots, and roots (Fig. 1) using both a weighted (bacteria F_1, 63_ = 12.5, R^2^ = 0.28, p < 0.001; fungi F_1, 57_ = 11.73, R^2^ = 0.29, p < 0.001) and an unweighted UniFrac distance matrix (bacteria F_1, 63_ = 7.75, R^2^ = 0.19, p < 0.001; fungi F_1, 57_ = 7.03, R^2^ = 0.21, p < 0.001). For bacterial and fungal communities, post-hoc contrasts showed differences between the structures of microbiomes in the inocula, shoots, and roots (p = 0.001 for all pairwise comparisons, FDR-corrected). Post-hoc tests highlighted a difference in the multivariate structure between root and shoot bacterial communities (UniFrac, p = 0.04; weighted UniFrac, p = 0.01; FDR-corrected) but not for fungal communities (UniFrac p = 0.81, weighted UniFrac p = 0.68, FDR-corrected).

**Figure 1.**
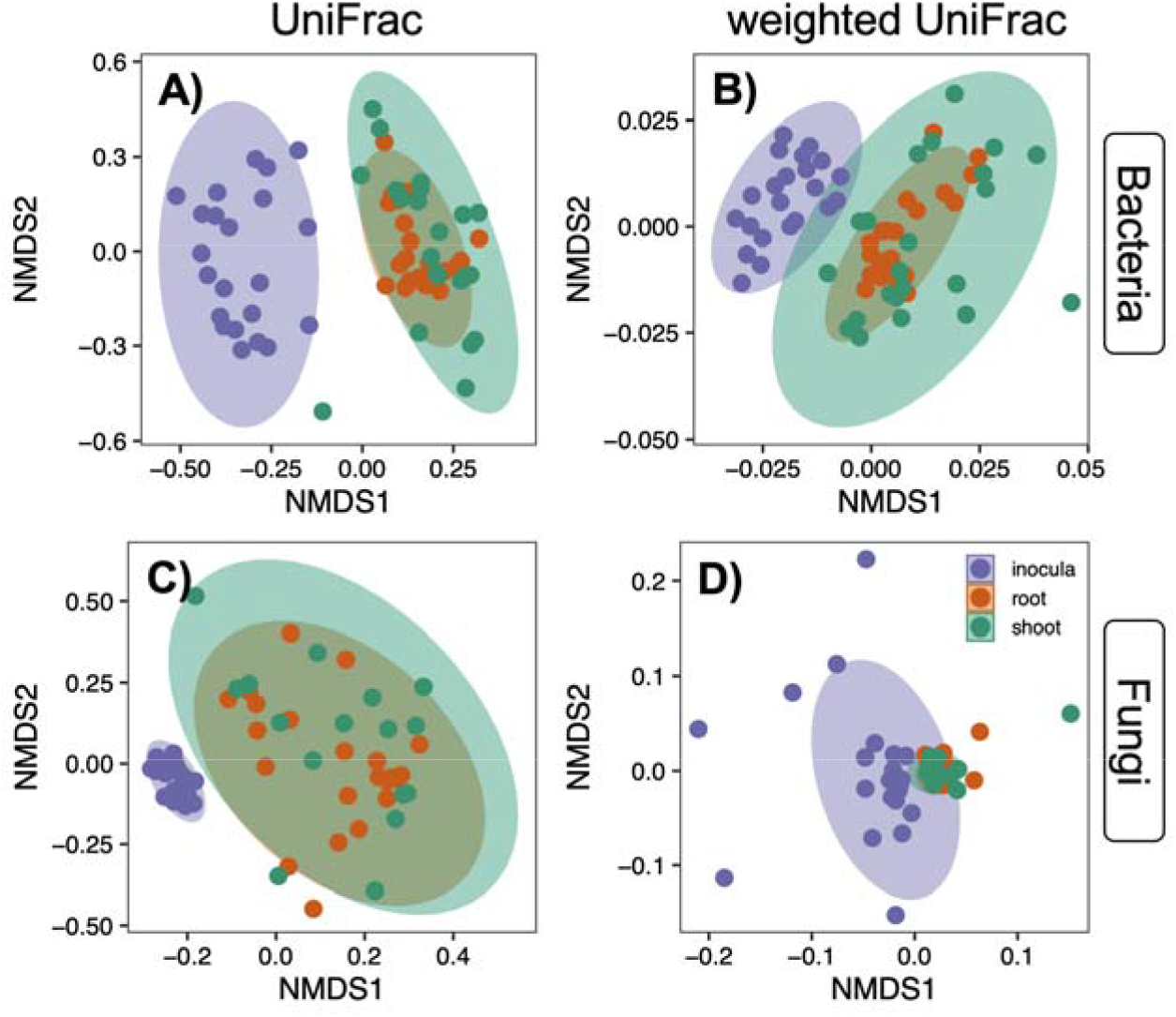
NMDS (Non-Metric Multi Dimensional Scaling) of bacterial (A, B) and fungal (C, D) communities of inocula, root, and shoot samples, using unweighted (A, C) and weighted (B, D) distances between samples. Ellipses represent the 95% CI for each compartment.

When focusing on the diversity of bacterial microbial communities, we found differences in phylogenetic diversity (F_2, 63_ = 282.21, p < 0.0001, Fig. 2A), Shannon’s diversity index (F_2, 63_ = 27.42, p < 0.0001, Fig. 2B), Simpson’s diversity index (F_2, 63_ = 9.16, p = 0.0003, Fig. 2C), and observed richness (F_2, 63_ = 33.76, p < 0.0001, Fig. 2D). Post-hoc contrasts showed higher phylogenetic diversity (Fig. 2A), Shannon diversity (Fig. 2B), and observed richness (Fig. 2D) in the inocula and roots than in the shoots, and a higher dominance in the shoots than in the other two compartments (Fig. 2C). Similarly, in the fungal community, we also found differences in phylogenetic diversity (F_2, 57_ = 505.41, p < 0.0001, Fig. 2E), Shannon’s diversity index (F_2, 57_ = 280.79, p < 0.0001, Fig. 2F), Simpson’s diversity index (F_2, 57_ = 38.97, p < 0.0001, Fig. 2G), and observed richness (F_2, 63_ = 498.44, p < 0.0001, Fig. 2H). Both roots and shoots showed lower microbial diversity and richness (Fig. 2E, F, H) and higher dominance (Fig. 2G) than the inocula, but there was no difference between the two plant compartments (Fig. 2E-H).

**Figure 2.**
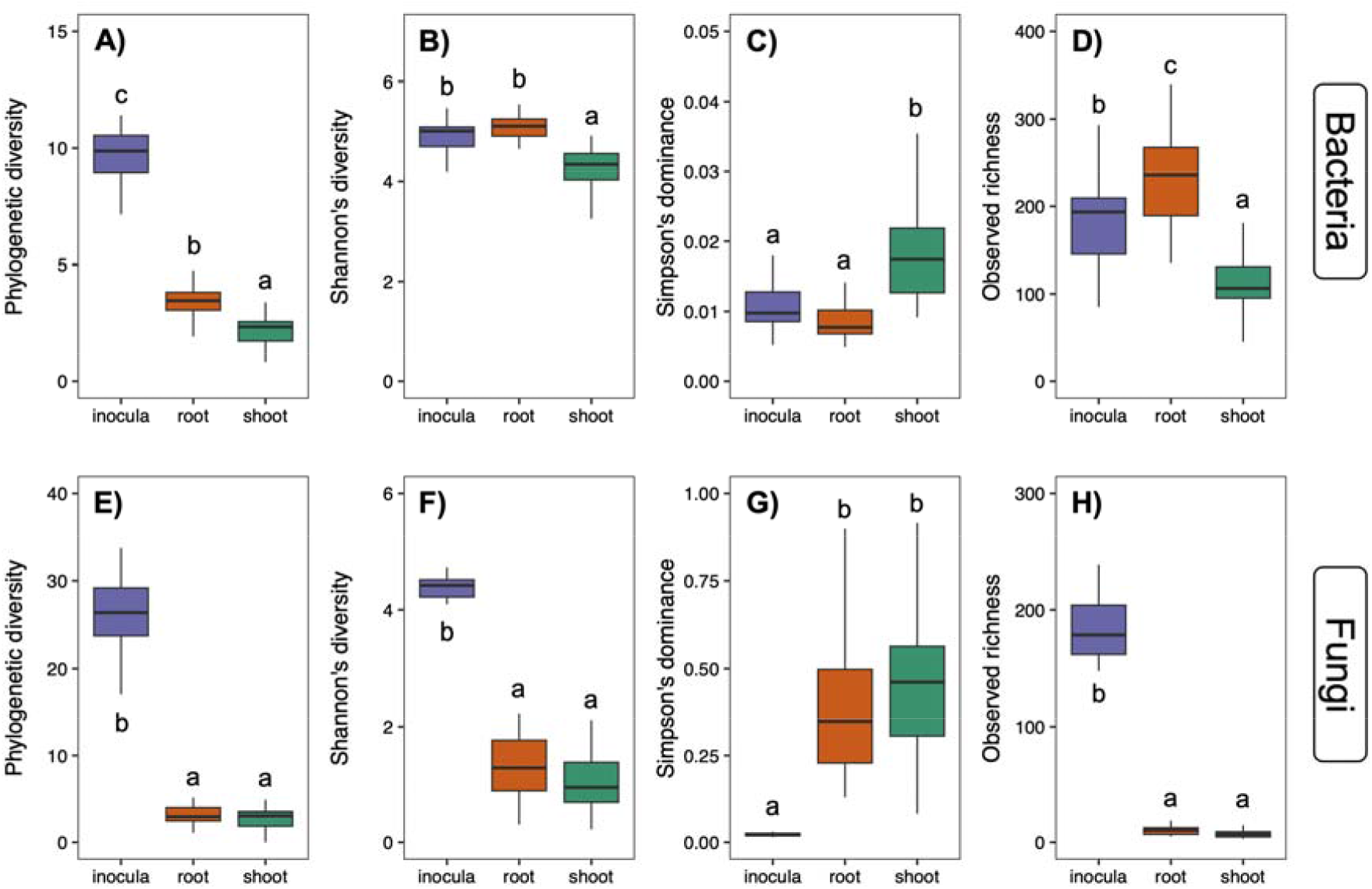
Phylogenetic diversity (A and E), Shannon diversity (B and F), Simpson dominance (C and G), and observed richness (D and H) indexes for bacterial (A, B, C, D) and fungal (E, F, G, H) communities in samples collected from inocula, root, and shoot. Pairwise comparisons are shown as letters for each boxplot, and exact p-values are reported in Tab. S1.

To test the hypothesis that the root microbiome is assembled from the inoculum and that the shoot microbiome is further selected from the root microbiome, we identified changes in ASV abundance between pairs of compartments. We found that 336 bacterial ASVs were enriched in the roots compared to the inocula, and 137 ASVs were enriched in the shoots compared to the inoculum (Fig. 3A and 3 B). Although 87 bacterial ASVs were enriched in both roots and shoots compared to the inoculum, no ASV was significantly enriched in the shoots compared to the roots (Fig. 3C). In contrast, only one fungal ASV was significantly enriched in the roots compared to the inoculum (Fig. 3D), and no ASV was enriched in the shoots compared to the inoculum (Fig. 3E) or roots (Fig. 3F).

**Figure 3.**
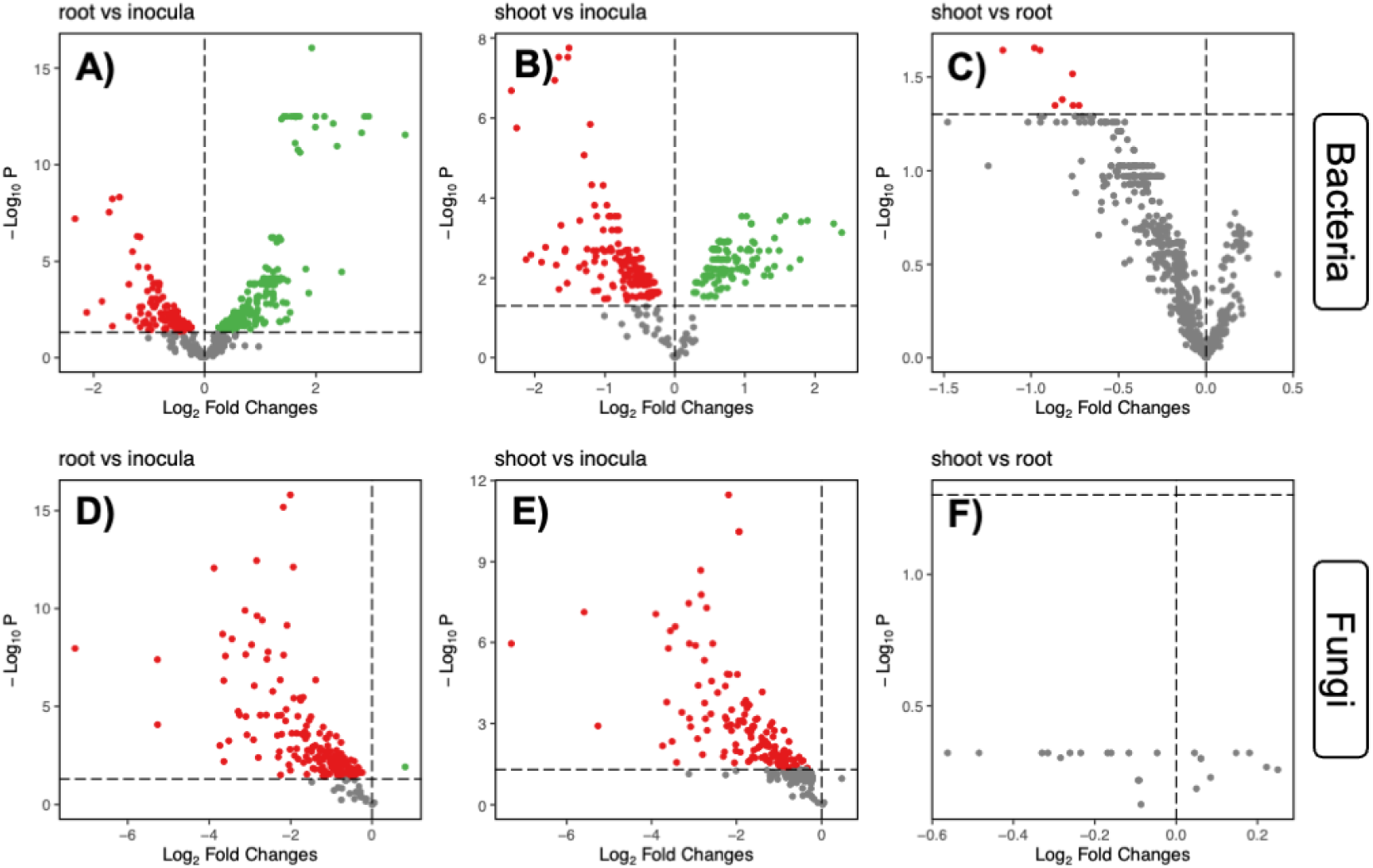
Volcano plots showing differentially abundant ASVs of bacteria (top) and fungi (bottom) between pairs of compartments. (A and D) root (green) vs inocula (red), (B and E) shoot (green) vs inocula (red), (C and F) shoot (green) vs roots (red). ASVs in grey are not differentially abundant between the two compartments.

Further tests showed that the number of ASVs shared between compartments (Fig. 4A and 4 B) was always different from random chance, except for fungal ASVs that were shared between all compartments (Fig. 4B). While the number of ASVs shared between compartments was lower than random chance in most cases, the number of bacterial ASVs shared between roots and shoots was higher than random (Fig. 4A). In addition, the number of bacterial and fungal ASVs unique to shoots was always lower than that of simulated random microbial communities, whereas the number of ASVs unique to the inoculum was always higher than random chance (Fig. 4).

**Figure 4.**
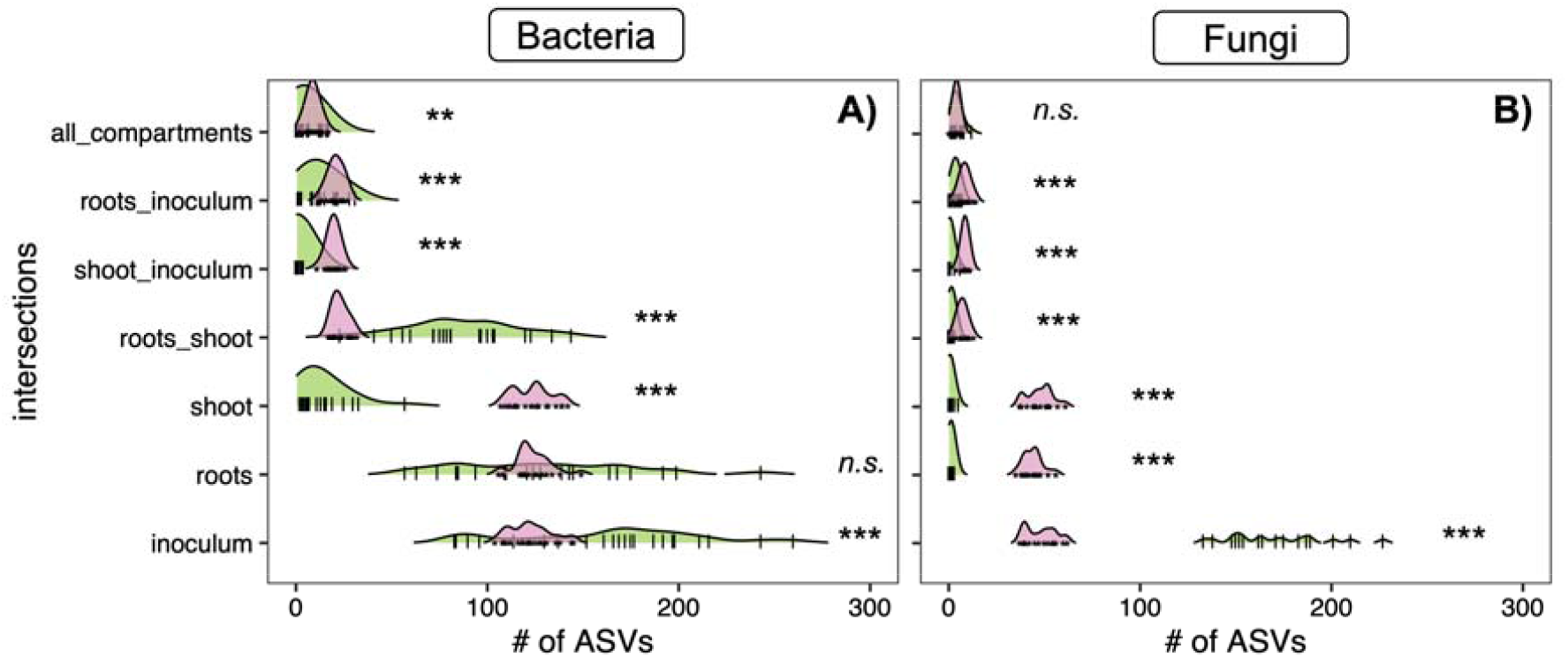
Ridgeline plot showing the distribution of the number of ASVs shared between compartments, and testing the difference between observed data (green distribution, vertical lines are individual datapoints) and randomized data (pink distribution, small stars are individual datapoints). Asterisks on the side of each pair of ridgelines show the results from a *lmer* testing for differences between the means of the two distributions (*** p<0.001, *n*.*s*. p > 0.05).

To further explore the idea that the microbiome of gnotobiotic plants is assembled through deterministic processes rather than stochastic associations from the inoculated microbial communities, we tested the assembly processes of the microbial communities associated with plants that were not inoculated, thus interacting only with microbes that were inherited from the seeds. Thus, when examining the composition of the microbial communities associated with plants that had not been inoculated, we observed a community composed of 172 bacterial and 9 fungal ASVs (Fig. S4). When examining the composition of these communities, we found that the composition was highly variable between samples, suggesting that microbiome assembly under gnotobiotic conditions followed stochastic rules. We further tested this idea by calculating the βNTI index for both bacterial and fungal communities and found that the βNTI of both root and shoot microbiomes was between -2 and 2 (Fig. 5A and 5 B), suggesting that stochastic processes are the major driver of microbiome assembly in plants associated with microorganisms derived solely from seeds. In addition, the RCbray index for roots was on average 0.35 (bacteria) and 0.04 (fungi), while for shoot was on average 0.28 (bacteria) and -0.05 (fungi). We then examined plants that had been inoculated with microorganisms from the field and found that the βNTI index was always > 2 (Fig. 5C and 5D), suggesting that deterministic processes contributed to the assembly of plant microbiomes.

**Figure 5.**
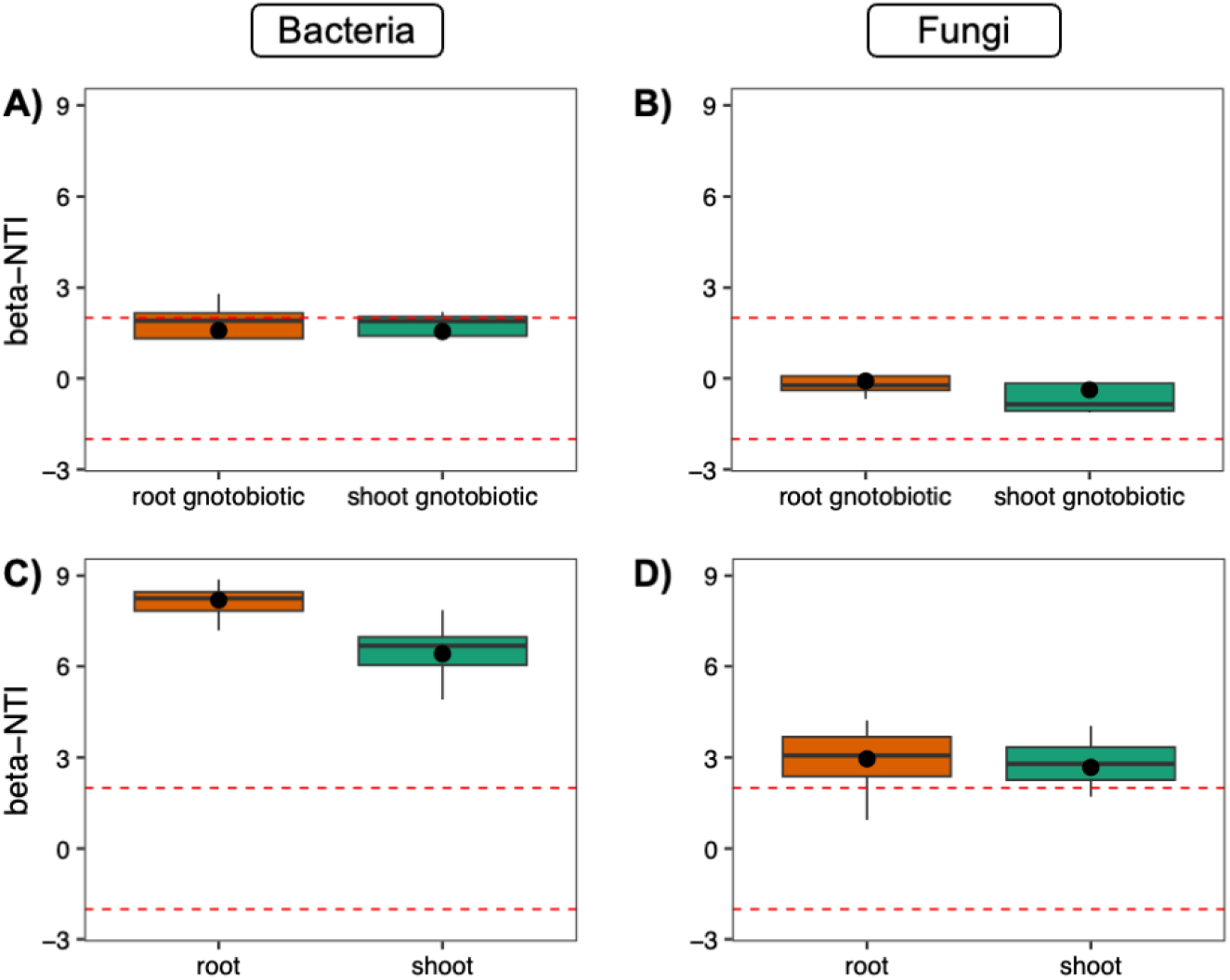
βNTI (beta-nearest taxon index) of bacterial (A and C) and fungal (B and D) communities associated with gnotobiotic (A and B) and inoculated (C and D) lettuce plants. Horizontal dashed lines represent reference values of -2 and 2, indicating that the thresholds in the microbiome assembly are considered to be shaped by stochastic processes. The black dot at the top of the boxplot represents the distribution mean.

## Discussion

In this study, we showed that gnotobiotic plants can quickly select inoculated microbial communities and assemble root and shoot microbial communities via deterministic processes. First, our results showed that the structure of the microbial communities associated with the inoculated plants was different from that of the inoculum, and that the microbial diversity was higher in the roots than in the shoots. This result suggests that the plants created a selective environment in the root and shoot compartments. We observed a high proportion of ASVs with differences in abundance between roots/shoots and the inoculum, whereas no ASV was significantly enriched in the shoot compared to the roots. This suggests that plants exert a selective force on root-associated microbial communities, and from this, they are able to assemble the shoot microbiome. Finally, we directly tested the prevalence of deterministic or stochastic assembly processes in both the gnotobiotic and inoculated plants. Our results suggest that microbial communities in gnotobiotic plants, which are built from microbial taxa inherited from seeds, are not driven by selection processes but by ecological drift (RCbray index < 0.95) [28]. However, once inoculated, plants were able to quickly (one week) assemble root and shoot microbiomes through deterministic processes. Taken together, our results support our hypothesis that seedlings create a selective environment belowground and that through selection from the soil microbial community, they assemble both root and shoot microbiomes. In addition, we found support for our hypothesis that the inherited seed microbiome is subjected to stochastic assembly processes, whereas, once exposed to the environmental microbiome (i.e., inocula in our case), plants can exert selective pressure and assemble their microbial communities through deterministic processes.

When testing the assembly processes of bacterial and fungal communities in maize under field conditions and across developmental stages, Xiong et al. [14] found that plant bacterial communities were assembled through selection at early growth stages. Similar results were observed in the wetlands of *Typha orientalis* [22], where the microbiome of seedlings showed signatures of selection rather than stochasticity in their assembly. In both cases, samples were collected from the field, and while this ensured that the results hold in real-life conditions, these observations might be biased by external factors that might influence the assembly processes of the plant microbiome. In the present study, we used a reductionist approach to grow gnotobiotic plants and exposed them to a range of inocula, thus removing possible interference from the air microbiome and abiotic effects. Thus, we were able to distinguish the selection effect driven by the plant from other possible factors driving selection on the soil microbiome. Most previous studies have taken a snapshot of a particular stage of plant development, while Xiong et al. [14] followed maize plants throughout development and found that the assembly of plant bacterial communities was dominated by deterministic processes early in development, while stochastic processes were more dominant later in the growth season. This matches our results in that plants at early stages exert a stronger selection on their microbiome. While more data needs to be gathered, this might support the idea that plants vary the strength of the selection they impose on their microbiome across development, in the same way they redirect resources (e.g., root exudates) at more mature developmental stages [29].

Several other studies have focused on understanding the ecological processes driving the assembly of the plant microbiome, and as reported in the introduction, the results vary greatly across plant phylogeny, geography, and plant organs. This variation may be caused by several factors (e.g., plant genotype and stressors) that are known to influence the plant microbiome [1] and are difficult to account for in field settings. For example, differences in the strength of selection have been observed across rice genotypes [15] or in response to biotic and abiotic stressors [19, 25]. However, these patterns appear to be conserved among closely related species. Wang et al. [21] and Yan et al. [23] found that stochastic processes drive the assembly of microbial communities associated with *Eucalyptus* plants. Similarly, Guo et al. [12] and Yin et al. [15] observed deterministic processes that guided the assembly of rice microbiomes. Our study was performed under heavily controlled conditions; thus, it would be useful to disentangle the plant-driven effect from other factors that might bias the results. However, this information needs to be paired with experiments under field conditions, where other factors might mask the microbiome selection driven by the host plant. This is a key step towards disentangling the effects of the host plant within the holobiont.

Our results also show that the deterministic processes driving the assembly of shoot and root microbial communities in our system are dominated by variable selection (βNTI index > 2). Observing a βNTI index lower than 2 would indicate that plants will assemble similar microbial communities starting from different inocula (homogenous selection). This observation, although interesting, might simply be generated by the use of different inocula, which generate more dissimilar plant-associated microbial communities. This is not a surprising result, as it is widely acknowledged that plants growing in soils hosting different microbial communities are associated with dissimilar microbial communities. On the other hand, plants are continuously challenged by growing on soils hosting very dissimilar microbial communities, including spatial variation across small (seed dispersal) and large scales (seed commercialization), temporal variation at small (within the same year) or large scales (across multiple years, e.g., soil seed banks), plant-soil feedbacks, plant range expansion, and several other events. Although plants might be associated with different microbial taxa in different contexts, they might still exert selective pressure on function rather than identity, as microbial communities with different structures might code for similar functions [30]. Several studies have suggested that plants can modulate their microbiome through changes in root exudates and VOCs [31–33], including the selection of specific microbial functions that might differ throughout plant development [31]. This mechanism might explain why the same plant genotype grown in association with different microbial communities might have little phenotypic variation. This is an interesting perspective that needs to be addressed by multi-omics studies that go beyond the taxonomic composition of the microbial community and focus on its functional role in the host plant. In addition to the idea that the selection of dissimilar microbiomes might be entirely driven by the host plant, we also need to consider that different soil microbial communities might be characterized by different interactions between the members of the microbiome, which in turn might influence the final outcome of plant-microbiome interactions [34].

Interestingly, we did not observe any differences in the structure of the microbial communities between the shoots and roots. This contrasts with the general idea that plant-associated microbial communities are mainly assembled by organ or compartment [34, 35], and there is evidence that this differentiation can also be detected in gnotobiotic conditions [35, 36]. However, our study focused on seedlings at the early developmental stage, whereas previous studies mainly focused on plants at a later developmental stage. This might explain the inconsistency of the results, suggesting that the shoot microbiome is first assembled from the soil/root microbiome, but then differentiated through microbial recruitment or transient association with microorganisms from the environment. Future studies may provide more detailed evidence for this idea. In addition, when focusing on gnotobiotic plants not exposed to inocula, we observed a community composed of 172 bacterial and nine fungal ASVs that were highly variable at the level of individual seedlings. This is consistent with results from previous studies on the microbiome of single seeds, which showed a very high variability in the composition of the microbial community between individual seeds [37, 38].

Our study contributes to expanding our understanding of the mechanisms that guide the assembly of the plant microbiome, suggesting that plants can drive selection processes at early developmental stages. Although this idea needs to be tested with a wider set of hosts and under different conditions, it provides further evidence that will help clarify patterns in the assembly of microbial communities across plant species. This information is key to understanding the functioning of the plant microbiome and how we can direct its assembly to influence the host or guide us in the assembly of synthetic microbial communities that can help achieve more sustainable agriculture and ecosystem restoration.

## Methods

### Experimental procedure

Lettuce seeds (variety “Romabella”) were surface-sterilized using the method of Davoudpour et al. [39], with a few modifications. Briefly, lettuce seeds were treated with 70% ethanol for 3 min before being sterilized twice with a 30 mL mixture of 8% sodium hypochlorite and 17 μL Tween 20 (15 min each round, 30 min in total). The seeds were thoroughly rinsed five times with sterile water. Sterilized seeds were then placed on wet filter paper in a sterile Petri dish for approximately ten days under direct sunlight for germination. Seed sterility was checked by placing 15 seeds on PDA medium and incubating them for approximately 5 days at 20°C, after which no microbial growth was observed. Surface sterilization was performed to remove external contaminants that could rapidly grow in sterilized soil and influence plant growth.

The experiment was performed using sterile microboxes (Combiness Europe, Nevele, Belgium; 14 cm H × 9 cm base Ø, 1 L volume) commonly used for micropropagation, allowing plant growth under sterile conditions. Each microbox was sterilized for 10 min with 4% sodium hypochlorite before being autoclaved at 121°C for 15 min, filled with approximately 170 g of autoclaved soil, and then watered with 10 ml of autoclaved water. The autoclaved soil was prepared by sieving the soil to 1 mm to remove large particles, which were watered, covered, and left for approximately seven days at ∼20°C to allow the growth and development of microorganisms. After that, the soil was autoclaved at 121°C for 3 h, allowed to cool to room temperature for approximately 24 h, and autoclaved again at 121°C for 3 h before being used to fill the sterile microboxes.

Five seedlings (∼1 week old) were transplanted into each microbox and inoculated with 1 mL of different soil inocula (see below). Each microbox was inoculated with a different inoculum (*n* = 21). In addition, four microboxes were inoculated with 1 mL of distilled water as a control. Soil inocula and sterile water were added to the soil to avoid direct contact with seedlings. After that, all boxes were kept under direct sunlight at room temperature (24-25°C) and rearranged every 24h to account for variation in light exposure. Seven days after soil inoculation, all the plants in each microbox were gently collected using their entire root system. Plants were rinsed with autoclaved water to remove soil particles before being dissected into two parts (shoots and roots) and placed separately in 2 ml Eppendorf tubes, pooling all five seedlings from the same microbox. Subsequently, all samples were freeze-dried for 24 h and then crushed for 1 min at 30 Hz using a bead mill homogenizer and 2-3 glass beads (3 mm Ø). Finally, the samples were stored at -80°C until DNA extraction.

### Soil inocula

Soils were collected from different areas with varying cultivation methods, crops, forests, and uncultivated land to obtain a diverse microbial community between each inoculum. Each soil inoculum was prepared according to the method described by Walsh et al. [40] with minor modifications. Briefly, 20 g of soil was transferred to a 50 mL sterile falcon tube filled with 20 mL of sterile distilled water. The tubes were then vortexed for approximately 30 s before being centrifuged at 1000 ×*g* for 1 min to sediment the larger soil particles. The supernatant was transferred to a new Falcon tube and the microbes were pelleted by centrifugation at 3200 ×*g* for 5 min. The supernatant was discarded, and the pellets were resuspended in 20 mL of sterile phosphate-buffered saline (PBS). A small aliquot of each inoculum was stored at -80°C for further processing.

### DNA extraction, amplicon library preparation and sequencing

DNA from the roots and shoots was extracted using the PowerPlant® Pro DNA Isolation Kit (MO BIO), whereas DNA from the inocula was extracted using the PowerSoil® DNA Isolation Kit (MO BIO), according to the manufacturer’s instructions. The DNA concentration and quality were estimated using a NanoDrop 2000 spectrophotometer (Thermo Fisher Scientific). Bacterial communities were characterized by amplifying a portion of the 16S rRNA gene using the primers 515f/806r [41], while fungal communities were characterized by amplifying the ITS2 region using the primer pair ITS3-KYO2/ITS4 [42]. Amplifications were performed using standard two-step PCR, first amplifying the target fragment and then ligating the adaptors/barcodes for sequencing [see, for example, our previous study 43]. Amplifications included: (i) non-template controls (*n* = 3), where DNA extraction was performed by replacing samples with nuclease-free water to account for possible contamination of instruments, reagents, and consumables used for DNA extraction; and (ii) negative PCR controls (*n* = 3), in which the DNA template for PCR was replaced with the same volume of ultrapure water. Libraries were then quantified using a Qubit fluorometer (Thermo Fisher Scientific), pooled together at equimolar ratios, and sequenced on an Illumina MiSeq platform (Illumina, CA, USA) using the MiSeq Reagent Kit v3 600 cyclers (300PE) following the supplier’s instructions.

### Data analysis

Data analysis was performed using R v4.1.2 [44] and visualizations were created using *ggplot2* [45]. Paired-end reads were processed using the *DADA2* pipeline [46] to remove low-quality data, identify Amplicon Sequence Variants (ASVs) and remove chimeras. Taxonomy was assigned using SILVA v138 [47] for bacteria, and UNITE v8.2 [48] for fungi. The ASV table, metadata, taxonomical annotation for each ASV, and phylogenetic tree of all ASVs were merged into a *phyloseq* object [49] for handling. Before downstream analyses, all ASVs identified as “chloroplast” or “mitochondria” were discarded, and the package *decontam* [50] was used to remove potential contaminants using data from non-template and negative controls (see above). ASV sequences were aligned using *DECIPHER* [51] and bootstrapped maximum-likelihood phylogenetic trees were estimated using *phangorn* [52]. After removing singletons, the ASV table was normalized using the package *wrench* [53] and used for all analyses, except when calculating the diversity metrics (observed richness and Faith’s, Shannon’s, and Simpson’s indices), which were integer numbers that needed to be used. All analyses were performed separately for bacterial and fungal communities.

Differences in the multivariate structure of microbial communities between the three compartments (inocula, roots, and shoots) were tested using PERMANOVA (999 permutations) on both a weighted UniFrac and an unweighted UniFrac distance matrix between samples, performed using the package *vegan* [54]. Results were visualized using a NMDS (non-metric Multi-Dimensional scaling) procedure, and pairwise contrasts were inferred using the package *RVAideMemoire* [55], correcting p-values using the FDR (False Discovery Rate (FDR) procedure.

The diversity of microbial communities within each sample was estimated using Faith’s phylogenetic diversity index calculated using the package *picante* [56], and the observed richness, Shannon’s diversity, and Simpson’s dominance indices were calculated using the package *microbiome* [57]. Differences between compartments (inocula, roots, shoot) were tested using the packages *lme4* [58] and *car* [59] by fitting a separate linear model for each diversity index and using “compartment” as fixed factor. Pairwise contrasts were inferred using the package *emmeans* [60], correcting p-values using the False Discovery Rate (FDR) procedure.

ASVs that were differentially abundant between pairs of compartments (inocula, roots, and shoots) were identified using the package *MaAsLin2* [61], using an adjusted p-value of 0.05.

We also calculated the number of ASVs shared between pairs of compartments and all three compartments for each inoculum, and repeated this procedure by randomizing the values within the ASV table. We then tested for differences between the two distributions (observed vs. random number of shared ASVs between compartments) using a generalized linear mixed-effects model, with category (observed, random) as a fixed effect and the inoculum ID as a random variable.

The beta Nearest Taxon Index (βNTI, quantifying the deviation of Mean Nearest Taxon Distances from null expectations) was calculated using the package *picante* and used to test whether microbial communities assembled following deterministic or stochastic assembly processes [62, 63]. The RCbray index was estimated using the package *iCAMP* [28].

## Acknowledgements

This work has been supported in by the Italian Ministry of Education, University and Research (MIUR) under the 2017 Program for Research Projects of National Interest (PRIN) with Grant n. 20172TZHYX “A gnotobiotic-based approach to unravel the role of the plant microbiome and develop synthetic communities increasing plant growth and stress tolerance.”

## Authors contribution

Conceptualization: NM, AM, and LS; Methodology: NM and AM; Investigation: NM and AM; Visualization: NM and AM; Funding acquisition: LS; Writing-original draft: NM; Writing-review and editing: all co-authors.

## Data availability

Raw sequencing data are available on the NCBI SRA under Bioprojects PRJNA1029922 (16S) and PRJNA1029924 (ITS).

## Code availability

Data and code to replicate analyses are available at: https://github.com/amalacrino/gnotobiotic_lettuce

