## Supplementary material for "Lettuce seedlings rapidly assemble their microbiome from the environment through deterministic processes"

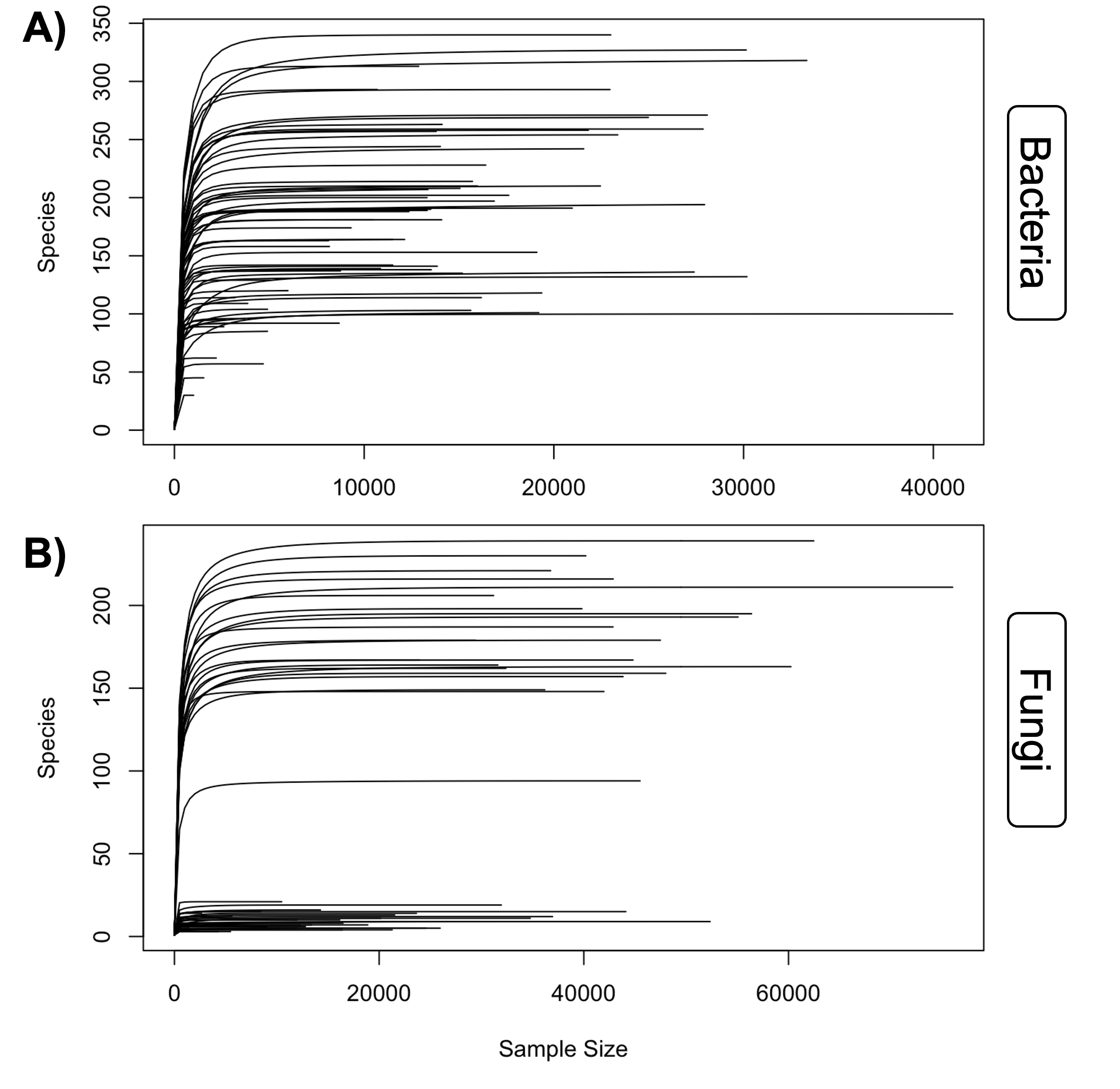

**Figure S1**. Rarefaction curves for (**A**) 16S and (**B**) ITS datasets after cleanup and removal of plastidial reads.

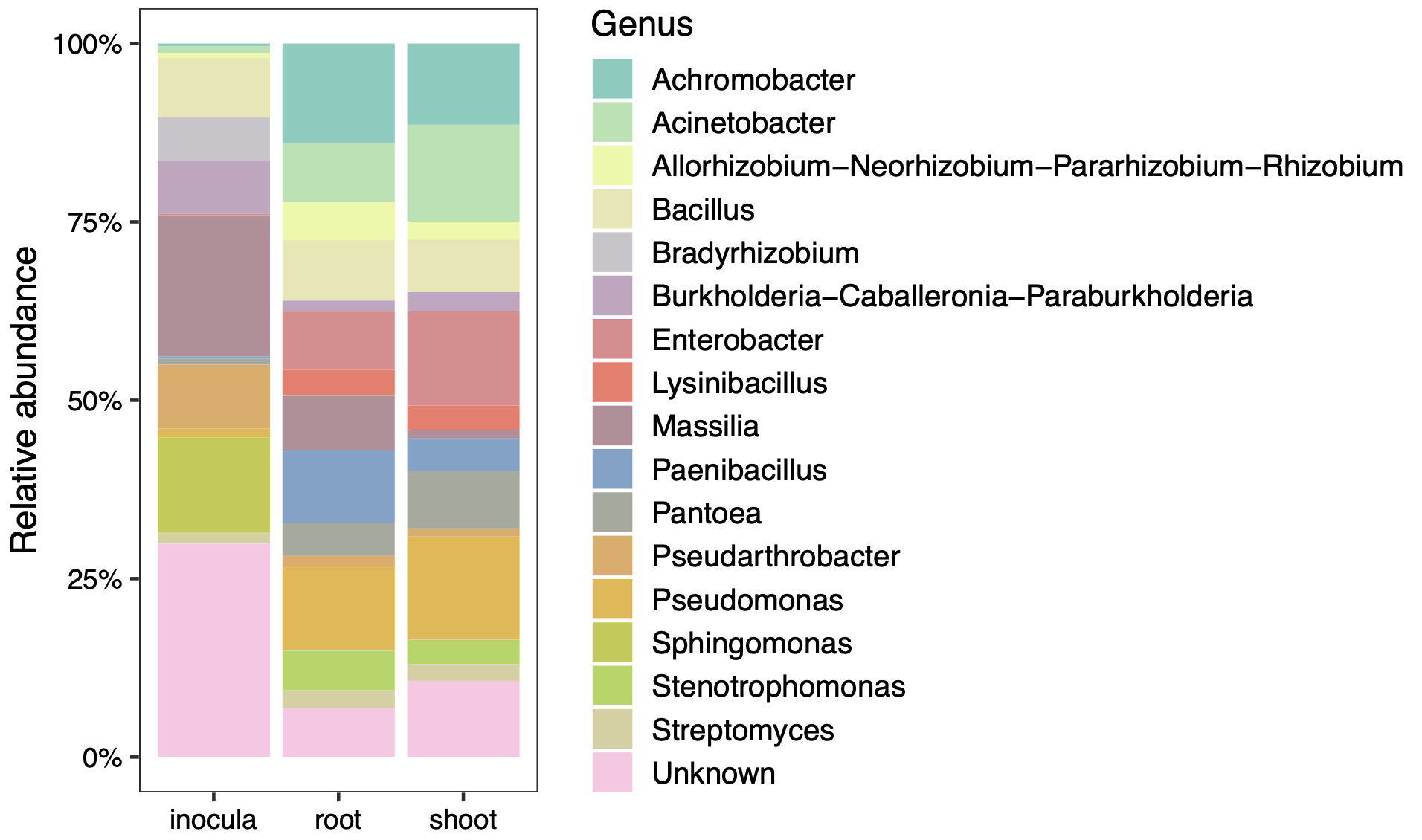

**Figure S2**. Relative abundance (%) of bacterial taxa in inoculated lettuce plants.

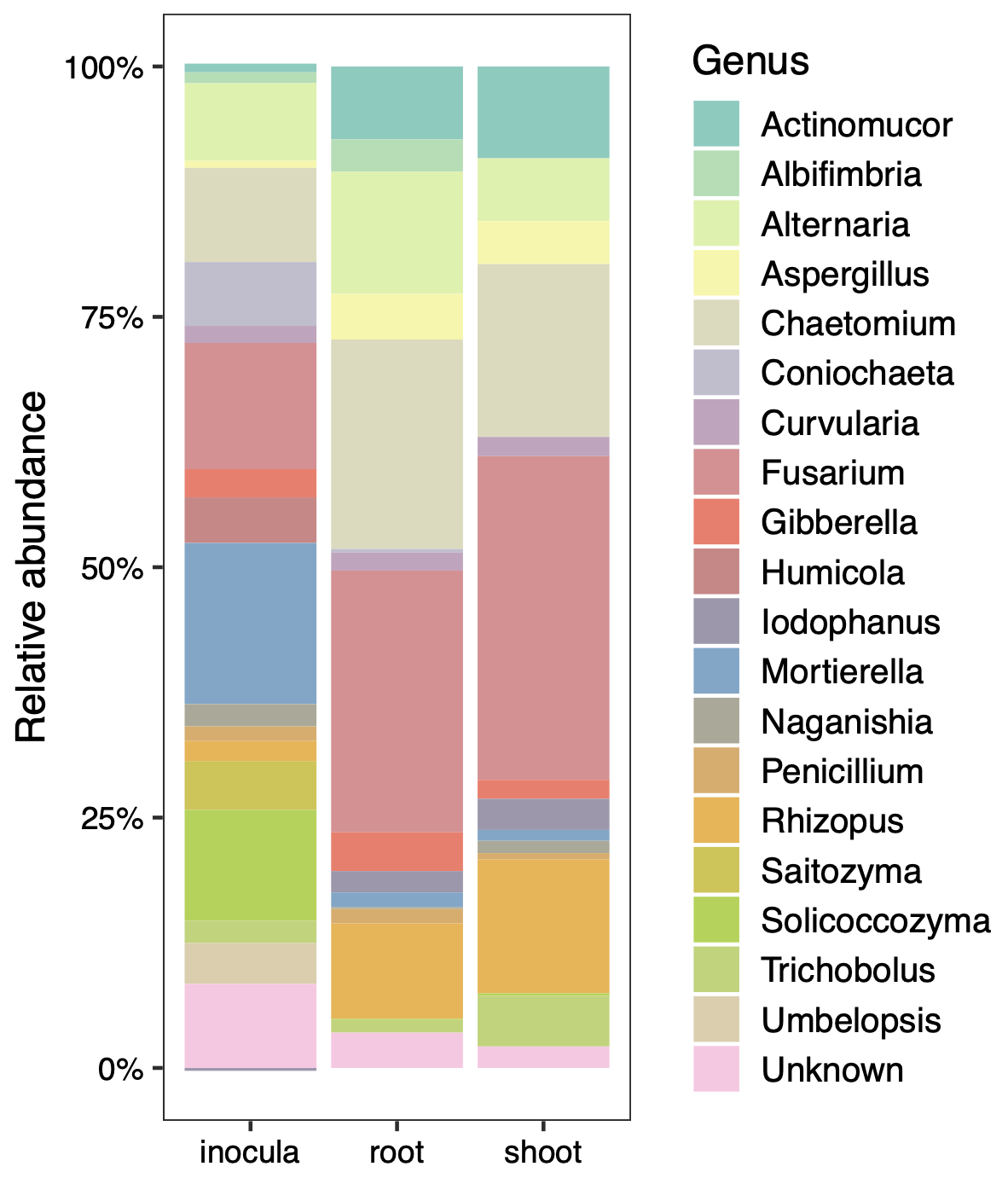

**Figure S3**. Relative abundance (%) of fungal taxa in inoculated lettuce plants.

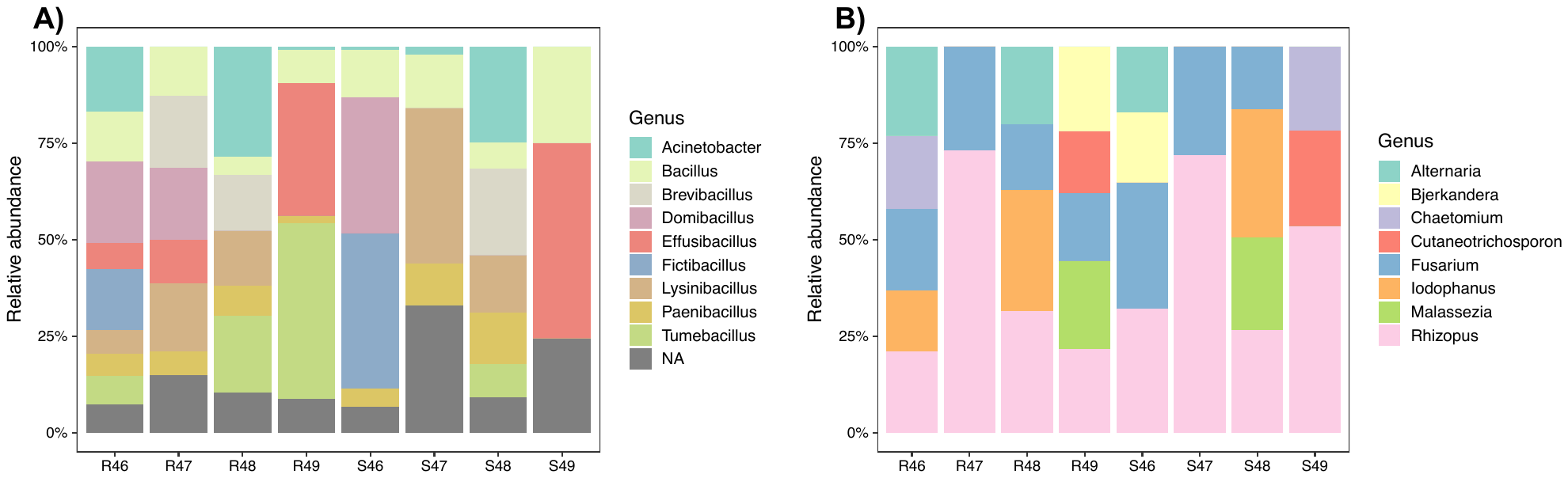

**Figure S4**. Relative abundance (%) of (**A**) bacterial and (**B**) fungal taxa in gnotobiotic lettuce plants. Samples starting with the letter “R” indicate root samples, while those starting with the letter “S” indicate shoot samples.

**Table S1**. Pairwise comparisons between compartments for different diversity indices, for both bacterial and fungal communities.

| **Community** | **Diversity index** | **Comparison** | **p-value (FDR)** |
| --- | --- | --- | --- |
| Bacteria | Phylogenetic diversity | inocula – root | <0.001 |
|  |  | inocula – shoot | <0.001 |
|  |  | root – shoot | <0.001 |
|  | Shannon’s diversity | inocula – root | 0.167 |
|  |  | inocula – shoot | <0.001 |
|  |  | root – shoot | <0.001 |
|  | Simpson’s dominance | inocula – root | 0.362 |
|  |  | inocula – shoot | 0.017 |
|  |  | root – shoot | <0.001 |
|  | Observed richness | inocula – root | 0.004 |
|  |  | inocula – shoot | <0.001 |
|  |  | root – shoot | <0.001 |
| Fungi | Phylogenetic diversity | inocula – root | <0.001 |
|  |  | inocula – shoot | <0.001 |
|  |  | root – shoot | 0.815 |
|  | Shannon’s diversity | inocula – root | <0.001 |
|  |  | inocula – shoot | <0.001 |
|  |  | root – shoot | 0.512 |
|  | Simpson’s dominance | inocula – root | <0.001 |
|  |  | inocula – shoot | <0.001 |
|  |  | root – shoot | 0.440 |
|  | Observed richness | inocula – root | <0.001 |
|  |  | inocula – shoot | <0.001 |
|  |  | root – shoot | 0.927 |
